# Emergence and diversification of a highly invasive chestnut pathogen lineage across south-eastern Europe

**DOI:** 10.1101/2020.02.15.950170

**Authors:** Lea Stauber, Thomas Badet, Simone Prospero, Daniel Croll

## Abstract

Invasive microbial species constitute a major threat to biodiversity, agricultural production and human health. Invasions are often dominated by one or a small number of genotypes, yet the underlying factors driving invasions are poorly understood. The chestnut blight fungus *Cryphonectria parasitica* first decimated the American chestnut and a recent outbreak threatens European chestnut trees. To unravel the mechanisms underpinning the invasion of south-eastern Europe, we sequenced 188 genomes of predominantly European strains. Genotypes outside of the invasion zone showed high levels of diversity with evidence for frequent and ongoing recombination. The invasive lineage emerged from the highly diverse European genotype pool rather than a secondary introduction from Asia. The expansion across south-eastern Europe was mostly clonal and is dominated by a single mating type suggesting a fitness advantage of asexual reproduction. Our findings show how an intermediary, highly diverse bridgehead population gave rise to an invasive, largely clonally expanding pathogen.

**Data availability:** All raw sequencing data is available on the NCBI Short Read Archive (BioProject PRJNA604575)

## Introduction

Over the past century, a multitude of invasive species have emerged as threat to forest and agricultural ecosystems worldwide (Wingfield et al., 2010; Santini et al., 2013). Increased human activities, such as global trade, enabled invasive species to cause economic damage through reduced agricultural production, degradation of ecosystems and negative impacts on human health (Marbuah et al., 2014). A key group of invasive species in forests are fungal pathogens, which are often accidentally spread via living plants and plant products (Rossman, 2001). To successfully colonize a new environment, fungal pathogens have to overcome several invasion barriers including effective dispersal abilities, changes in available hosts, competition with other fungi, and niche availability (Gladieux et al., 2015). This may be achieved with plastic phenotypic changes followed by rapid genetic adaptation (Garbelotto et al., 2015). However, because the number of initial founders is often low, invasive populations are frequently of low genetic diversity which reduces adaptive genetic variation (Allendorf & Lundquist, 2003; Yang et al., 2012). Yet, many fungal plant pathogen invasions were successful despite low genetic diversity within founding populations (Raboin et al., 2007; Fontaine et al., 2013; Wuest et al., 2017).

A major hypothesis explaining the successful expansion of invasive populations despite low initial genetic diversity is the so-called bridgehead effect (Lombaert et al., 2010). In this model, highly adapted lineages emerge through recombination among genotypes established in an area of first introduction. Hence, the primary introduction serves as the bridgehead for a secondary and more expansive invasion. Although the bridgehead effect has been proposed as a scenario for many biological invasions (Gau et al., 2013; van Boheemen et al., 2017), empirical evidence for the creation of highly adaptive genotypes within bridgehead populations is still largely missing (Bertelsmeier & Keller, 2018). An alternative explanation suggests that first introductions simply serve as repeated sources of inoculum for additional secondary invasions without creating adaptive genotypes *in situ* (Bertelsmeier & Keller, 2018). This alternative scenario implies that initial populations are already composed of genotypes adapted to the new environment or have high phenotypic plasticity (Bock et al., 2015; Gladieux et al., 2015; Vuković et al., 2019). Dissecting whether invasive species were pre-adapted to the new environment or gained adaptation through a bridgehead effect is crucial for effective containment strategies, but requires a deep sampling of genotypes during the early invasion process.

The pace of adaptive evolution leading to successful invasions is determined by several life-history traits including the mating system (*i*.*e*. sexual, asexual or mixed). The low availability of mating partners can be a major cost for individuals at the invasion front. To avoid mating costs, invasive fungal pathogens often switch from sexual to predominantly asexual reproduction (Heitman et al., 2013; Suehs et al., 2004). However, the switch to asexual progeny reproduction limits genetic diversification and adaptive potential (Taylor et al., 2015; Drenth et al., 2019). Populations of some highly successful invaders, such as the ascomycete *Ophiostoma novo-ulmi* (Paoletti et al., 2006), or the oomycete *Phytophthora ramorum* (Grünwald et al., 2012), are dominated by a single mating type. Low diversity in invasive lineages may be breached by secondary invasions introducing the opposite mating type as observed in *Phytophthora infestans* in Europe. Prior to the 1980s only the A1 mating type was present followed by the secondary introduction of the A2 mating type (Mariette et al., 2016). Introgression from closely related species can also reintroduce a missing mating type. *O. novo-ulmi* acquired the missing mating type from *O. ulmi* (Brasier & Webber, 2019; Paoletti et al., 2006). Although switches in reproductive modes can be a key factor for invasion success (Philibert et al., 2011), mechanisms underlying such switches remain poorly understood (Billiard et al., 2012).

The ascomycete *Cryphonectria parasitica* (Murr.) Barr. is the causal agent of chestnut blight, a lethal bark disease of *Castanea* species (Rigling & Prospero, 2018). The pathogen is native to eastern Asia and was first observed in 1904 in North America on the American chestnut (*Castanea dentata*). In the following years, the disease rapidly spread throughout the distribution range of *C. dentata*, causing the vast decimation of this native tree species (Elliott & Swank, 2008). In 1938, the fungus was first detected on the European chestnut (*C. sativa*) near Genoa (Italy) and is now established in all major chestnut-growing countries in Europe (Rigling & Prospero, 2018). The damage to the European chestnut may have been reduced by the presence of the *Cryphonectria hypovirus 1* (CHV-1) which acts as a biological control agent of chestnut blight (Rigling & Prospero, 2018). The virus can be transmitted both vertically to asexual spores (conidia) or horizontally through hyphal anastomoses between virus-infected and virus-free strains (Heiniger & Rigling, 1994). Hyphal anastomoses are controlled by a vegetative compatibility system and the virus spreads most effectively between strains of the same vegetative compatibility type (Cortesi et al., 2001).

Population genetic analyses showed that the invasion of Europe occurred through multiple introductions both from native populations in Asia and from bridgehead populations in North America (Dutech et al., 2012). Invasive European *C. parasitica* populations are characterized by lower vegetative compatibility diversity than North American and Asian populations (Milgroom & Cortesi, 1999). Furthermore, European populations exhibit lower recombination rates (Dutech et al., 2010; González-Varela et al., 2011; Prospero & Rigling, 2012). Mating in *C. parasitica* is controlled by a single mating type (MAT) locus (Marra & Milgroom, 2001) and natural populations can reproduce both sexually and asexually (Marra et al., 2004). Previous analyses of south-eastern European *C. parasitica* populations based on sequence characterized amplified region (SCAR) markers suggested that the region was largely colonized by a single and likely asexual lineage also identified as S12 (Milgroom et al., 2008). The lineage belongs to the vegetative compatibility type EU-12. Within the distribution range of the lineage, sexual structures (*i*.*e*. perithecia) are rarely found (Sotirovski et al., 2004; Milgroom et al., 2008). Based on SCAR marker and field records, Milgroom et al. (2008) suggested that the invasive S12 lineage originated in northern Italy and a subsequently spread across south-eastern Europe (Avolio, 1978; Biraghi, 1946; Buccianti & Feliciani, 1966; Karadžić et al., 2019; Myteberi et al., 2013; Robin & Heiniger, 2001). However, due to the low molecular marker resolution and challenges in relying on observational data, the origin, invasion route and genetic diversification of *C. parasitica* in south-eastern Europe remain largely unknown.

In this study, we sequenced complete genomes of a comprehensive collection of European *C. parasitica* strains with a fine-scaled sampling throughout the S12 invasion zone. Based on high-confidence genome-wide single nucleotide polymorphisms (SNPs), we identified the most likely origin and recapitulated the invasion process across south-eastern Europe. We show that the invasive *C. parasitica* lineage arose through an intermediary, highly diverse bridgehead population. During the expansion, the lineage became dominated by a single mating type but retained the ability to reproduce sexually.

## Results

### Genome-wide polymorphism analyses and phylogenomic reconstruction

We analyzed complete genomes of 188 *C. parasitica* isolates covering the European outbreak region of the invasive S12 lineage, as well as reference isolates from South Korea (*n* = 2) and the United States (*n* = 4). Isolates were sequenced at a mean depth of 8–27.5X to detect high-confidence genome-wide SNPs. A region of 179’501 -2’084’312 bp on scaffold 2 was associated to the mating type locus based on association mapping *p*-values (Supp. Fig. 2). This region encoded known mating type associated genes (Idnurm et al., 2015) and was characterized by a high SNP density consistent with observations in other fungi (Taylor et al., 2015). We removed SNPs within the mating type associated region to avoid confounding genetic structure with mating type divergence. We retained 17’873 SNPs and constructed a whole-genome maximum likelihood phylogenetic tree (Fig. 1A). The tree revealed two major clades splitting two Asian and one Swiss isolate (vegetative compatibility type EU-65) from the European and North American isolates. The large European/North American clade was subdivided into three clades. The clades a and c showed mixed geographic origins including North American isolates and European isolates of various vegetative compatibility (EU-)types (Fig. 1A, Supp. Fig. 3). In contrast, clade b consisted of genetically highly similar isolates (*n* = 113; Fig. 1A) of vegetative compatibility type EU-12, which predominantly originated from the Balkans, Italy, Turkey and Georgia. Approximately 93% (*n* = 105) of all isolates in the EU-12 clade were MAT-1 (Figure 1A) and were found in Albania, Bosnia, Bulgaria, Croatia, Georgia, Greece, Italy, Macedonia, Serbia and Turkey (Fig. 1B; Supp. Table 1). 6

**Figure 1:**
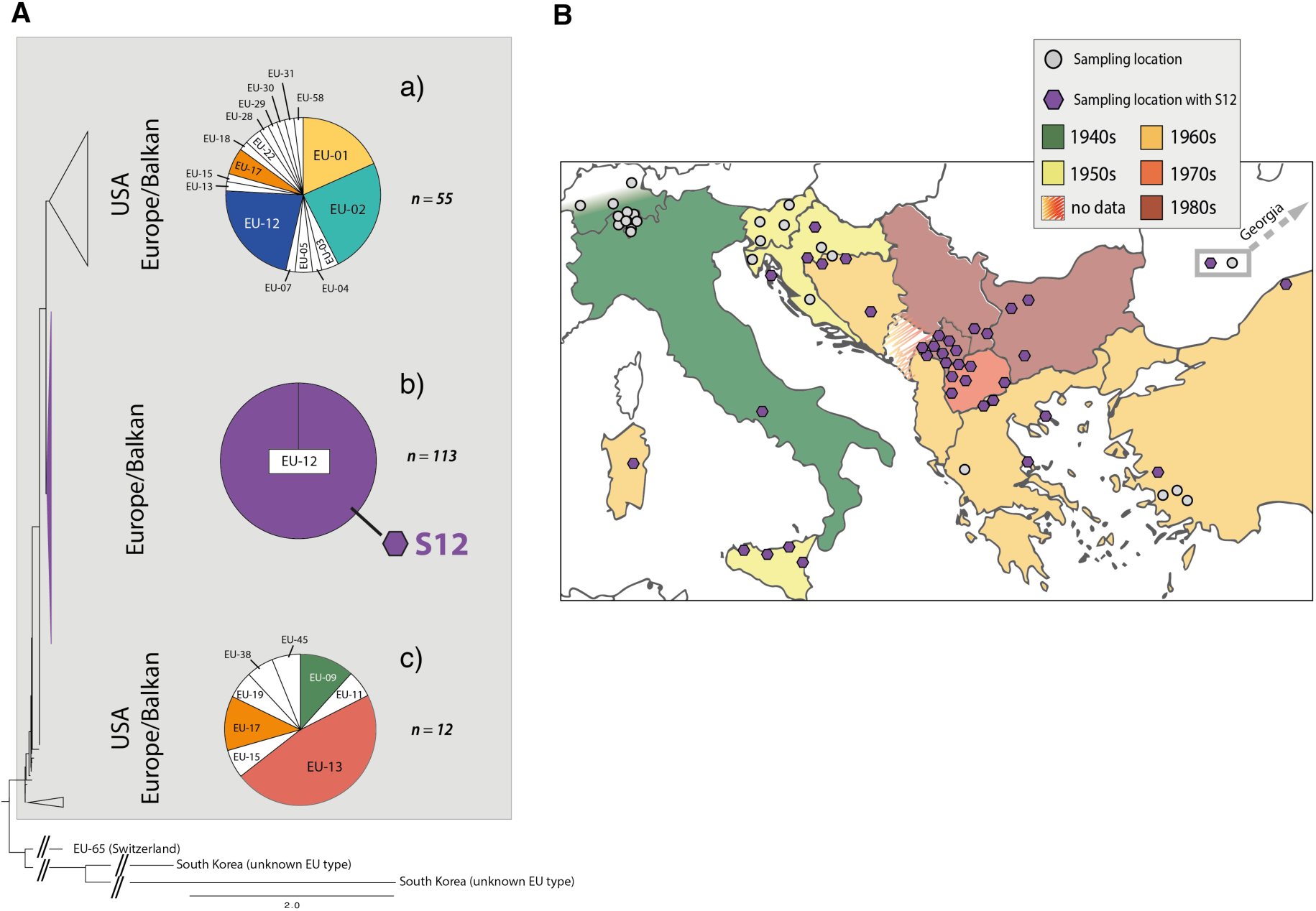
Genome-wide analyses of *Cryphonectria parasitica* isolates focusing on the invasion in south-eastern Europe. A) Whole-genome single nucleotide polymorphism (SNP) maximum likelihood tree constructed from 188 sequenced *C. parasitica* isolates. The US/European clade is shown with a grey box. The pie charts represent the proportions of vegetative compatibility types within each of the three clades. The most frequent vegetative compatibility types are highlighted in color (the color scheme matches with Figure 2). B) Map of European *C. parasitica* sampling locations. Grey circles show sampling locations of various vegetative compatibility types. Purple coloured hexagons represent sampling locations where EU-12 (mating type MAT-1) isolates were found (marked as “S12 lineage”). First observations of chestnut blight in the corresponding countries and regions are marked in a colour scheme according to decade. A tree with individual isolate labelling is shown in Supp. Fig. 3.

We performed a SplitsTree phylogenetic network analysis to account for reticulation caused by recombination. The network showed a high diversification, with both long branching and reticulation (Fig. 2, Supp. Fig. 4). The PHI-test revealed significant evidence for recombination (*p* < 0.0001). Despite the high level of genetic diversity, we found no evidence for geographic structure. Moreover, we found no clustering of isolates belonging to the same vegetative compatibility type with the exception of some EU-01, EU-02 and EU-12 (S12) genotypes from the Balkans. Nearly all *C. parasitica* isolates representing the S12 lineage showed almost identical genotypes and tight clustering. The most tightly clustered S12 genotypes were all of mating type MAT-1 (*n* = 104). Consistent with analyses by (Milgroom et al., 2008), this group represents the invasive S12 lineage at the origin of the expansion of *C. parasitica* across south-eastern Europe. Additionally, the phylogenetic network revealed closely related but not identical S12 genotypes of mating type MAT-2 (*n* = 7, Fig. 2). Hence, S12 outbreak strains of MAT-2 connect the nearly uniform cluster of S12 MAT-1 strains with the remaining genetic diversity of the major European subgroup of *C. parasitica*. The S12 cluster was furthermore connected with the remaining genotypes of the major clade by two EU-12 isolates from Bosnia (M1808 with MAT-1) and Georgia (MAK23 with MAT-2).

**Figure 2:**
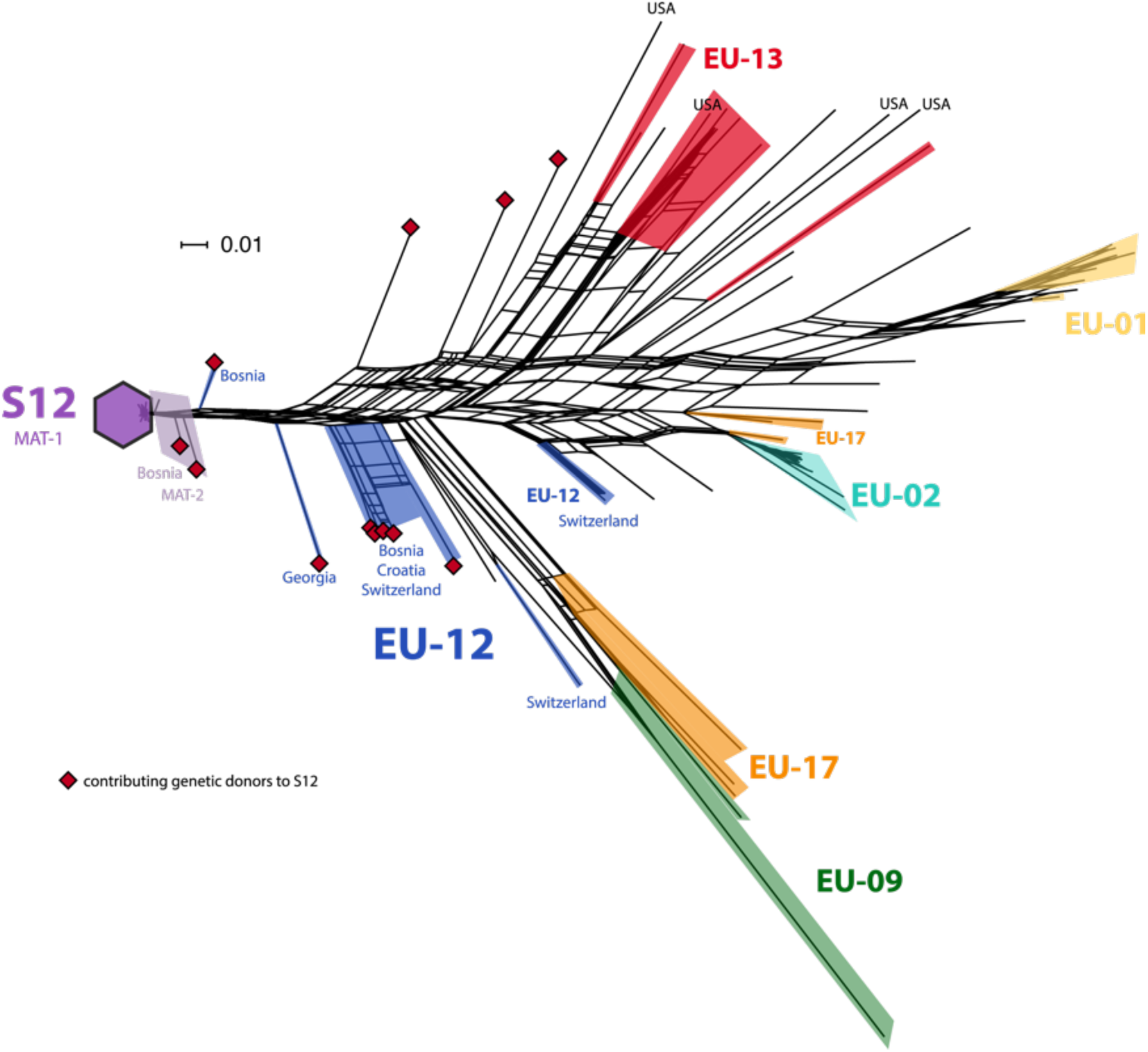
Phylogenetic network structure of the US/European subgroup analyzed by SplitsTree. The highlighted branches represent the most abundant vegetative compatibility types (color scheme matching Fig. 1A). Isolates belonging to the S12 outbreak lineage (EU-12; mating type MAT-1, *n* = 104) are marked with a purple hexagon. S12 isolates of mating type MAT-2 are highlighted in light-purple. Additional EU-12 isolates not belonging to the S12 lineage are highlighted in blue with information on the country of origin. Genetic donors of the S12 lineage as inferred by fineSTRUCTURE (Fig. 3, Table 1, Supp. Fig. 5) are marked with red squares.

### Identification of the most likely S12 founder populations

The maximum likelihood phylogenetic tree and the SplitsTree network revealed that the invasive S12 lineage has closely related genotypes occurring in Europe. Thus, to dissect the genetic origin of S12, we performed a co-ancestry matrix analysis using fineSTRUCTURE considering all isolates of the major North American/European subgroup, including the S12 lineage (*n* = 185; Figure 1A). The averaged co-ancestry matrix revealed no direct ancestors of the invasive S12 lineage among the major clade of different vegetative compatibility types. However, we found an association with a coefficient of 24.9– 32.3 between the recipient S12 genotypes and European donors from different locations (Fig. 3, Table 1, Supp. Fig. 5). The donors mostly originated from the North Balkans (Bosnia and Croatia), as well as Southern Switzerland, with the exception of isolate MAK23 from Georgia.

**Figure 3:**
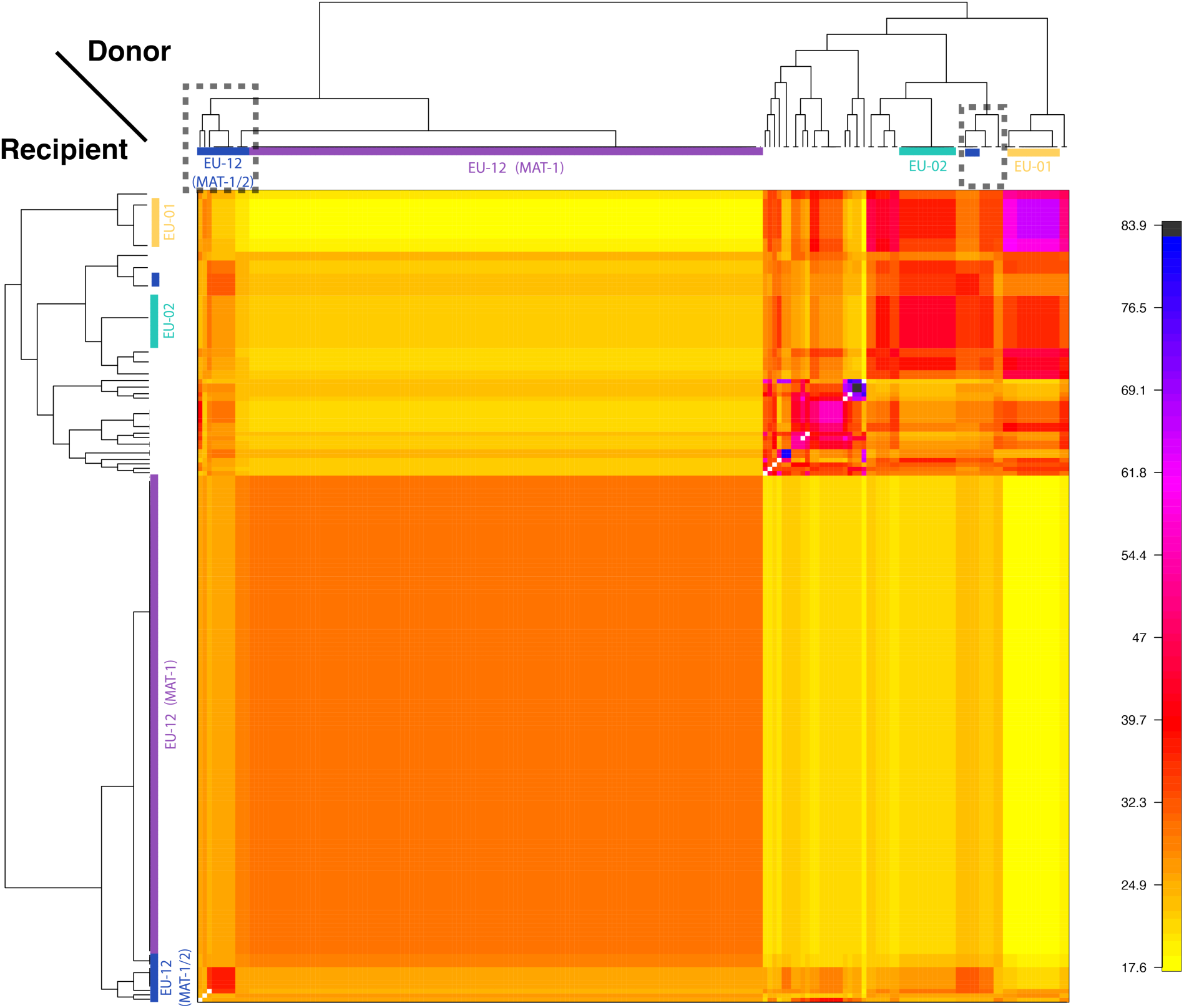
Analysis of donors to the S12 lineage genotypes. The averaged co-ancestry matrix and phylogenetic tree of the North American/European subgroup estimated by fineSTRUCTURE. The matrix shows recipient genotypes as rows and donors as columns. The heatmap indicates averaged coancestry coefficients between donors and recipients (i.e. darker colors indicate stronger genetic relationship between donors and recipients). Contributing S12 donors are marked in grey-dashed boxes. A detailed list of contributing donors is shown in Table 1, Fig. 2 and Supp. Fig. 5.

**Table 1:** Donor isolates for S12 identified using FineStructure. The geographic origin (country and population), vegetative compatibility (vc) type and mating type of donors contributing to genotypes of the invasive S12 lineage are given. See Fig. 3 for the corresponding co-ancestry matrix.

| Country | Population | vc type | Mating type | Donor isolate |
| --- | --- | --- | --- | --- |
| Bosnia | Projsa | EU-11 | MAT-1 | M1412 |
|  | Projsa | EU-12 | MAT-2 | M1407 |
|  | Projsa | EU-12 | MAT-2 | M1430 |
|  | Projsa | EU-12 | MAT-2 | M1431 |
|  | Vrnograč | EU-12 | MAT-1 | M1808 |
|  | Vrnograč | EU-15 | MAT-2 | M1834 |
|  | Vrnograč | EU-18 | MAT-2 | M1797 |
| Croatia | Kostajnica | EU-12 | MAT-1 | HK60B |
| Georgia | Mackhunceti | EU-12 | MAT-2 | MAK23 |
| Switzerland | Gnosca | EU-12 | MAT-1 | M6697 |
|  | Biasca | EU-12 | MAT-2 | M4023 |
|  | Biasca | EU-12 | MAT-2 | M4022 |
|  | Claro | EU-12 | MAT-2 | M2466 |

### Retracing S12 invasion routes and retention of mating competence

To infer potential invasion routes of the S12 outbreak lineage, we investigated intra-lineage genetic diversity across south-eastern Europe. We focused only on S12 isolates of mating type MAT-1 to delimit the closest genotypes contributing to the outbreak (*n* = 104; Fig. 2). The closely related genotypes segregated 468 high-confidence SNPs across the genome. The genetic structure assessed by a principal component analysis showed loose clustering of genotypes across south-eastern Europe (Fig. 4A and B, Supp. Fig. 6). We assigned genotypes to five regions: Italy, Northern Balkans, Central Balkans, Greece/Turkey and Georgia (Fig. 4A). Italy, Northern and Central Balkans, as well as Georgia harbored mainly genotypes of two dominant clusters. In contrast, the Greece/Turkey region contained genotypes of the two dominant clusters but also a broad diversity of further genotypes. We analyzed evidence for reticulation in the phylogenetic relationships among genotypes but found a star-like structure. We found minor evidence for reticulation among Central Balkans genotypes (Fig. 4C). Consistent with the phylogenetic network pattern, we found significant evidence for recombination within the S12 lineage (PHI test; *p* = 0.0035). We tested experimentally whether S12 mating type MAT-1 isolates were still able to reproduce sexually. We confirmed outcrossing of isolates of opposite mating type within the S12 lineage by pairing isolates from Molliq (Kosovo) and Nebrodi (southern Italy) (Fig. 4D). Mating pairs from Molliq and Nebrodi grew numerous perithecia, which are the fruiting bodies specific to sexual reproduction (Fig. 4E). Pairings of Bosnian isolates showed no perithecia formation. Using molecular mating type assays, we recovered both mating types among the ascospores produced from successful matings.

**Figure 4:**
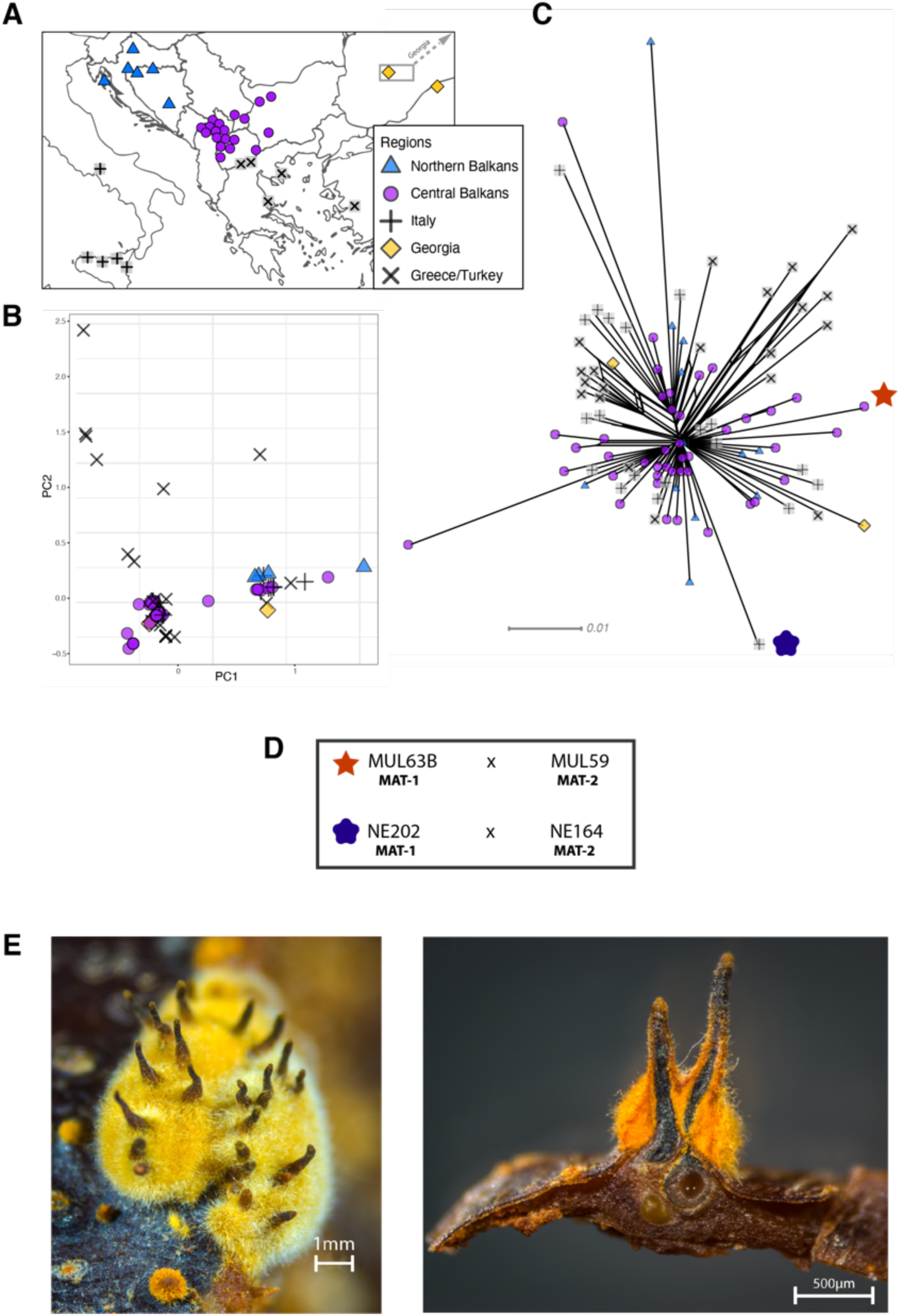
Fine-scale genetic diversity analyses of the S12 lineage. A) Geographic map showing the collected S12 isolates with mating type MAT-1 of *Cryphonectria parasitica* (*n*=104) according to five geographic regions. B) Principal component analysis (PCA) and C) SplitsTree of the S12 mating type MAT-1 outbreak isolates. Symbols and colors are as in A). D) Scheme of successful mating pairs of S12 mating type MAT-1 isolates crossed with isolates from the opposite mating type, of the same geographic origin. Symbols are as in C). E) Photographic images of sexual *C. parasitica* fruiting bodies (*i*.*e*. perithecia) emerging from crosses of S12 mating type MAT-1 isolates with isolates of the opposite mating type after five months of incubation under controlled conditions. Left: Perithecia embedded in a yellow-orange stromatic tissue. Right: Cross-section of perithecia and chestnut bark. Flask-shaped structures with a long cylindrical neck develop in yellow-orange stromatic tissue and are embedded in the bark (except for the upper part). The ascospores are formed in sac-like structures (asci) in the basal part of the perithecium. When mature, the ascospores are actively ejected into the air through a small opening (ostiole) at the end of the perithecial neck.

### Transposable element landscape and copy-number variation in the S12 lineage

Invasive pathogen lineages may have undergone crucial genomic rearrangements producing more fit genotypes. Here, we generated a *de novo* identification and annotation of transposable elements (TEs) for the *C. parasitica* genome. We found that 12% of the genome was composed of TEs with striking variation along the assembled scaffolds (*i*.*e*. quasi-chromosomes; Fig. 5A). In particular, regions on scaffold 2 matching the mating type locus are highly enriched in TEs suggesting that the large non-recombining region has undergone substantial degeneration (Fig. 5A). In contrast, the vegetative incompatibility (*vic*) loci are located in regions devoid of TEs. In fungal pathogens, effector genes and TEs are often co-localized in fast-evolving compartments of the so-called “two-speed genome” (Dong et al., 2015). However, the *C. parasitica* genome shows no apparent compartmentalization into gene-sparse and gene-rich regions. We used machine learning to predict secreted proteins most likely acting as effectors to manipulate the host. In contrast to other plant pathogens, effector gene candidates showed no tendency to localize in gene-sparse regions of the genome (Figure 5B). Non-repressed TEs can potentially create additional copies in the genome leading to intra-species variability in TE content. To detect such TE activity, we performed genome-wide scans of *C. parasitica* isolates for presence or absence of TEs based on split read and target site duplication information. At the loci with detectable TE presence/absence polymorphism, we found an over two-fold variation in total TE counts across the genome (Fig. 5C). The total TE count variation among the genetically diverse non-S12 isolates (North America and Europe only) was larger than the clonal S12 isolates. Nevertheless, the TE count variation among the S12 was surprisingly high given their recent emergence and extremely high similarity across the genome (Fig. 5C). This suggests that TE activity may have continued after the emergence of the S12 lineage and has created *de novo* genetic variation.

**Figure 5:**
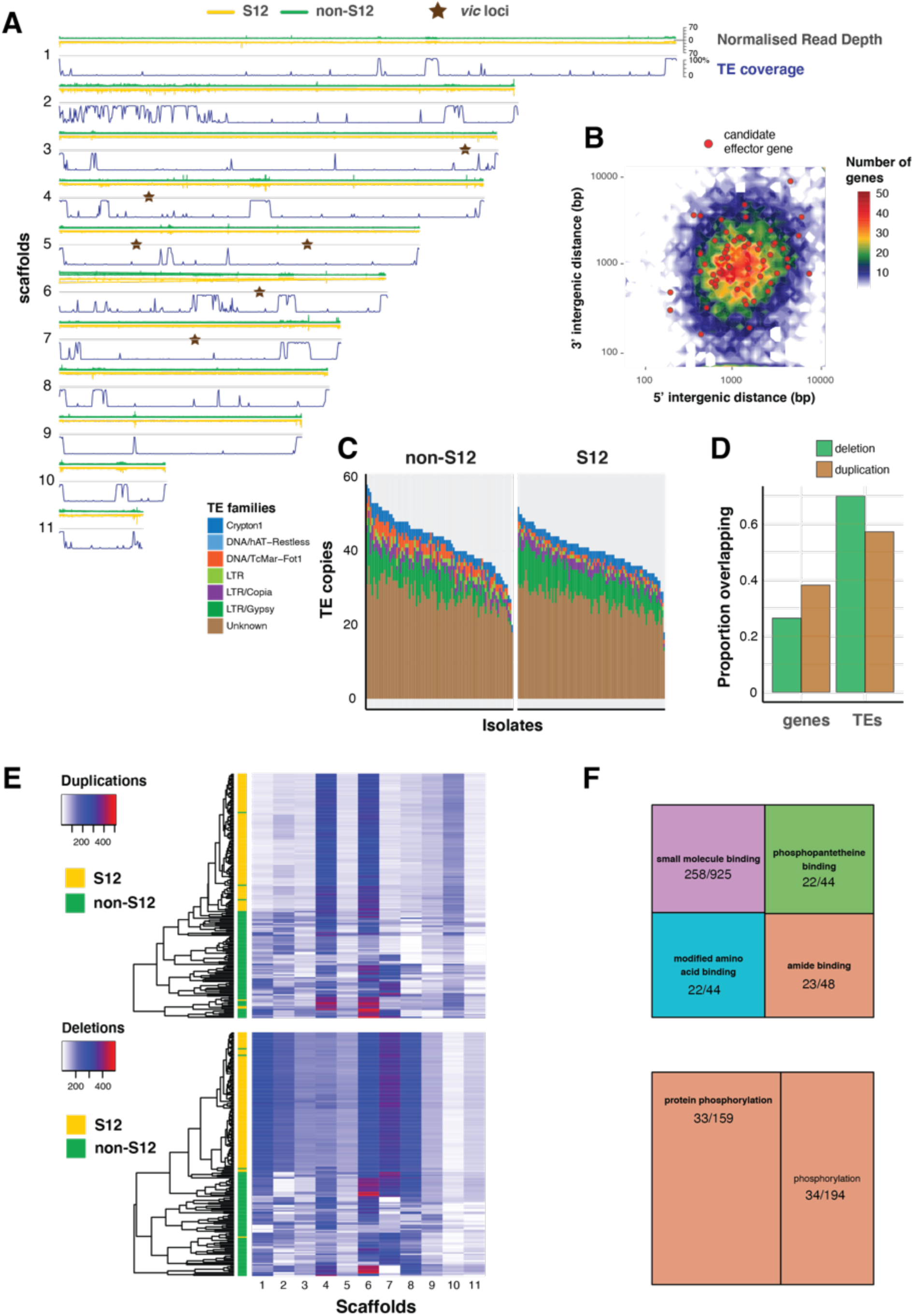
Transposable element (TE) landscape and copy-number variation. A) Genome-wide coverage of transposable elements (in 10kb windows) matched by with normalized read depth for the S12 and non-S12 lineages (North America and Europe only). *vic* loci: vegetative incompatibility loci.B) Genome-wide distribution of intergenic distances according to the length of 5’ and 3’ flanking regions. Red dots represent genes encoding predicted effector proteins. C) Counts of detected TE sequences across S12 and non-S12 isolates using split reads and target site duplication information. D) Proportion of normalized read depth windows (800 bp) with evidence for duplications (normalized read depth > 1.6) or deletions (< 0.4) overlapping with genes and TEs. E) Heatmap showing the number of windows (800 bp) with duplications and deletions. The dendrogram shows the similarity in duplication or deletion profiles for S12 and non-S12 isolates (North America and Europe only). F) Molecular functions (based on gene ontology) enriched in duplicated and deleted regions (upper and lower panel, respectively). Significance of the enrichment was at an alpha = 0.05 Bonferroni threshold. The numbers represent the number of genes with the matching gene ontology term in a duplicated or deleted region, and across the genome, respectively.

Repetitive sequences such as TEs can trigger non-homologous recombination leading to copy-number variation including deletions. We used normalized read coverage to assess copy-number variation across the species including the S12 lineage (Fig. 5A). Genes tend to overlap duplications rather than deletions and TEs tend to overlap deletions rather than duplications (Fig. 5D). The mating type region on scaffold 2 and the rDNA locus on scaffold 6 show particularly high levels of copy-number variation. Scaffolds 4 and 6 were overall rich in duplications and scaffolds 6 and 7 were rich in deletions (Fig. 5E). On average the S12 lineage bears more deletions per isolate compared to non-S12 lineages (∼71.2 versus 64.6 respectively). In a joint analysis of all isolates (except from Asia), we found that coding sequences overlapping with duplications and deletions are enriched for gene ontology terms associated with protein binding functions and protein phosphorylation activity, respectively (Fig. 5F). Overall, our results show that the S12 lineage underwent specific gene deletion and duplication patterns compared to the broad diversity of non-S12 isolates.

### Polymorphism and allele frequency spectra within the outbreak lineage

To gain insights into evolutionary forces shaping polymorphism in the outbreak S12 mating type MAT-1 lineage versus non-S12 populations, we first analyzed allele frequencies across the genome in both groups (Fig. 6A). The S12 lineage segregated virtually no intermediate allele frequencies in the range of 0.05–0.95. In contrast, the non-S12 genotypes showed overall a wide spectrum of allele frequencies across the genome. Genome-wide nucleotide diversity was extremely reduced in the S12 compared to non-S12 populations (Fig. 6B). We analyzed the predicted impact on protein functions of segregating mutations in the S12 lineage and non-S12 populations (Figure 6C). We found 4 highly and 94 moderately deleterious SNPs within S12 in contrast to 29 high and 773 moderately deleterious mutations in non-S12 groups (Fig. 6C). Three of the high impact SNPs in the S12 lineage were classified as stop gain mutations, as well as one splice acceptor variant (insertion variant). Two of these high impact mutations affect proteins of the major facilitator superfamily, as well as a protein containing a LCCL domain and an ecdysteroid kinase. Non-S12 populations showed an over-representation of low frequency high-impact mutations (Fig. 6C). This is consistent with purifying selection reducing the frequency of these mutations due to fitness costs. Within the S12 lineage nearly all segregating mutations were at very low frequency. We found only modifier (*i*.*e*. nearly neutral) mutations rising to higher frequency within the lineage. The extremely low level of polymorphism segregating within the S12 lineage prevents strong inferences of selective sweeps since the origin of the lineage. Hence, we focused on potential selective sweeps in the broader European and North American populations. To avoid a bias by the deep sampling of the S12 lineage, we excluded all but one of the S12 MAT-1 isolates (see Methods). We used RAiSD to produce a composite score of selective sweep signals and identified three strong outlier loci. The first sweep locus is located at the boundary of the mating type locus on scaffold 2 (Fig. 6D). The strongest sweep locus is on scaffold 3 and encompasses a ∼471 kb locus at positions 939-1’410 kb. The region contains 154 genes of which 107 encode conserved protein domains (Fig. 6D, Supp. Table 2). Ten genes overlap with SNPs showing the strongest signature of selection (*µ* statistic >40) of which two genes encode MFS transporters. The third selective sweep region of ∼190 kb region on scaffold 6 encompasses 60 genes (Fig. 6D, Suppl. Table S2). Three genes overlap with SNPs of *µ* > 30 and encode for a RTA1-like protein transporter, an oxidoreductase and an anaphase promoting protein, respectively (Fig. 6D, Suppl. Table S2).

**Figure 6:**
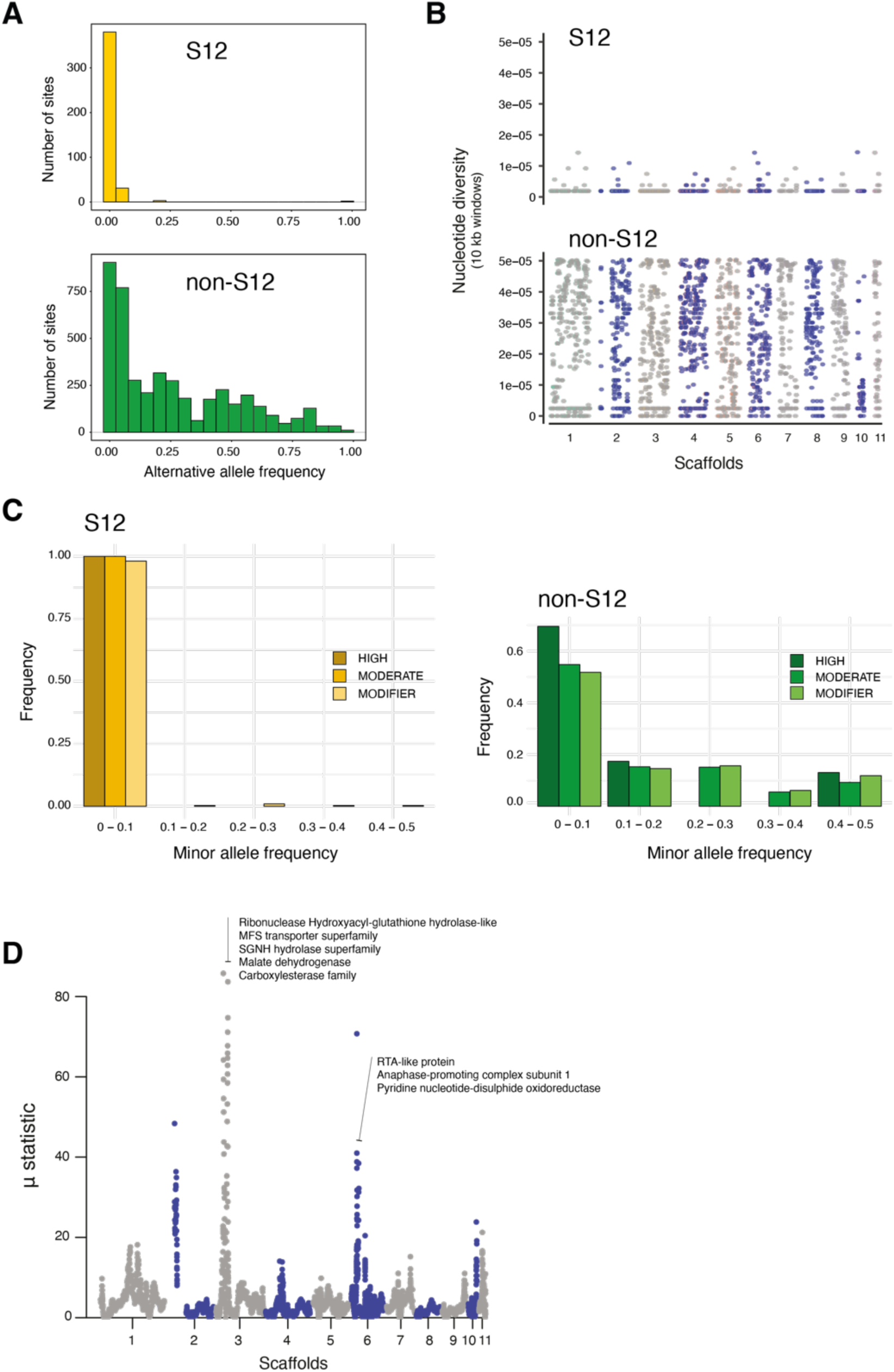
Polymorphism segregating in the S12 lineage. A) Alternative allele frequencies spectra across the genome for the S12 lineage (*n* = 104) compared to all other analyzed European (non-S12; *n* = 80). B) Genome-wide nucleotide diversity (pi) for the S12 lineage and for the non-S12 lineages in 10 kb windows. C) Minor allele frequency spectra of high, moderate and modifier (*i*.*e*. near neutral) impact mutations as identified by SnpEff. D) Genome-wide scan for selective sweeps (RAiSD). Encoded protein functions in the two top loci are shown as summaries.

## Discussion

We analyzed at the genome level how the chestnut blight pathogen *C. parasitica* successfully invaded south-eastern Europe after a first establishment on the continent. Deep sampling of the invasive genotypes showed that the invasion was caused by a single, highly homogeneous lineage consisting nearly exclusively of a single mating type. The invasive lineage likely transitioned recently from biparental mating populations to a single mating type. The spread across south-eastern Europe, suspected to originate in northern Italy, left no clear geographic imprint. This lack of genetic differentiation along the invasion route may be due to high levels of gene flow after the initial colonization or a very rapid spread from the center of origin to multiple locations across south-eastern Europe. Ongoing TE activity created unexpected levels of insertion polymorphism within the invasive lineage. The lineage carries a distinct gene deletion and duplication profile compared to the European and North American pool of *C. parasitica*. During the invasion, sexual reproduction was likely sporadically and may have introgressed genetic material from outside of the invasive lineage.

### The establishment of a European bridgehead population

Central and south-eastern European *C. parasitica* populations likely originated from North American lineages. Clusters of European and North American genotypes were largely overlapping indicating significant gene flow and, possibly, multiple introductions to Europe. These genome-scale analyses are consistent with previous findings documenting multiple North American introductions into France, but also directly from Asia (Milgroom et al., 1996; Dutech et al., 2012; Demené et al., 2019). Our results suggest that central and south-eastern European *C. parasitica* populations were largely established from North American sources alone. The high diversity at the genetic level with polymorphism at the level of SNPs, TEs and copy-number variation as well as at the level of vegetative compatibility types suggest that Europe was repeatedly colonized over the past century. Although genetic diversity could also have accumulated *in situ* in Europe, the establishment of a large set of genotypes and vegetative compatibility types seems difficult to explain with population age alone. Sexual recombination between the three most common vegetative compatibility types in Europe (*i*.*e*. EU-01, EU-02 and EU-05; Robin & Heiniger (2001)) could not account for the observed vegetative compatibility type diversity. Observational records date the first introductions into Europe to the 1930s (Robin & Heiniger, 2001). Hence, based on the observed genetic diversity, a scenario of repeated introductions of different vegetative compatibility types since 1930s seems most plausible.

Populations from Southern Switzerland, Slovenia Croatia and Bosnia are highly diverse, have nearly balanced mating type ratios, and a lack clustering according to vegetative compatibility types. This strongly suggests frequent outcrossing and population admixture, consistent with reports of perithecia in the field (Ježić et al., 2012; Prospero et al., 2006; Prospero & Rigling, 2012; Trestic et al., 2001). Low vegetative compatibility type diversity in most European *C. parasitica* populations was thought to have contributed to low population admixture within Europe compared to Asia and North America (Cortesi et al., 1996; Dutech et al., 2012; Prospero & Rigling, 2012). However, our genome-wide analyses revealed frequent and ongoing *in situ* admixture in Europe. Thus, vegetative compatibility type diversity does not necessarily underpin population admixture frequency and genetic diversity in sexually recombining populations. Our findings show that in asexually reproducing populations, such as in the S12 lineage, genotypes tend to cluster according to vegetative compatibility types.

### Emergence of an invasive lineage from a European bridgehead

The invasive lineage S12 most likely arose from existing genotypes established in Europe. We identified a series of closely related genotypes to the S12 lineage in Bosnia, Croatia, Georgia and southern Switzerland. Strikingly, the closest genotypes to the dominant S12 MAT-1 were S12 MAT-2 isolates found in Bosnia, Kosovo and southern Italy. Analyses based on a coancestry matrix identified a group of more distantly related genotypes from Bosnia, Croatia, Switzerland and Georgia having made the strongest genetic contributions to the S12 lineage. This shows that introductions from outside of Europe are unlikely to explain the emergence of S12. The lineage carries a unique set of copy-number variants compared to other European genotypes underlining the observation of a recombinant S12 genotype. Furthermore, the emergence of S12 was accompanied by a striking evolutionary transition from mixed mating type populations to single mating type outbreak populations. Human activity may have contributed to the shift towards single mating type populations. Shipments of infected chestnut seedlings from Northern Italy and other trading activities could have disseminated the invasive lineage further South. This would have exposed the pathogen to the geographically more fragmented chestnut forests typically found in south-eastern Europe where asexuality or selfing may be advantageous. Although *C. parasitica* is able to produce asexual conidia in large quantities, these specific spores are thought to be mainly splash dispersed by rain over short distances (Griffin, 1986). Accounting for occasional dispersal by birds or insects (Heald & Studhalter, 1914), conidia dispersal is unlikely to contribute substantially to the colonization of new areas.

Despite the loss of a mating type in the S12 lineage, we found genome-wide evidence for reticulation indicating at least low levels of recombination. If mating in *C. parasitica* follows the canonical process found in many ascomycetes, isolates of opposite mating type are required. Hence, S12 isolates of mating type MAT-1 may sporadically mate with rare S12 isolates of mating type MAT-2, which are comparatively more diverse. The emergence of the opposite mating type at low frequency could be the result of recombination with other genotypes and subsequent backcrossing. Combined with experimental evidence, we show that the dominant S12 mating type MAT-1 has retained the ability for sexual reproduction. Furthermore, in Bosnia, Croatia, Italy (Sicily) and Turkey the S12 lineage co-exists with other genotypes (*i*.*e*. vegetative compatibility types EU-01 and EU-02) of both mating types, potentially enabling sexual recombination and diversification *in situ*. The invasive S12 lineage was likely pre-adapted to the south-eastern European niche as we traced the origins to a likely Italian bridgehead population. Niche availability and benefits associated with asexual reproduction to colonize new areas may have pre-disposed the European *C. parasitica* bridgehead population to produce a highly invasive lineage.

### Expansion and mutation accumulation within the invasive S12 lineage

The MAT-1 S12 lineage diversified largely through mutation accumulation as nearly all high-confidence SNPs were identified as singletons. Mutation accumulation in absence of substantial recombination resulted in star-like phylogenetic relationships. We found a surprisingly high degree of differentiation among S12 genomes at the level of TE insertion polymorphism. Hence, active transposition of TEs is an important factor in diversifying the invasive lineage and possibly underpin future adaptive evolution. Analyses of allele frequency spectra suggested that the broader European *C. parasitica* populations efficiently removed the most deleterious mutations through purifying selection. In contrast, the S12 lineage shows strong skews towards very low minor allele frequencies of all mutation categories. Interestingly, we found a broader spread in allele frequencies for nearly neutral mutations in the S12 lineages. This suggests that despite the largely clonal population structure, deleterious mutations can still be removed through low levels of recombination and purifying selection. Using accumulated mutations as markers to retrace the spatial expansion of the invasive S12 lineage, we found no indication for a step-wise geographic expansion along potential invasion routes. A lack of genetic clustering across south-eastern Europe may be a consequence of high levels of gene flow frequently introducing new genotypes over large distances. However, the lack of geographic structure could also have its origins from substantial population bottlenecks during the spread of S12 across south-eastern Europe. Finally, the largely clonal lineage may also become exposed to processes such as Muller’s Ratchet fixing deleterious mutations over time (Felsenstein, 1974).

We show how a highly invasive fungal pathogen lineage can emerge from an intermediate, genetically diverse bridgehead population. This is in line with the self-reinforcement invasion model where initial introductions promote secondary spread (Bertelsmeier et al., 2018; Garnas et al., 2016). However, empirical evidence for adaptation in bridgehead populations is often elusive. Additionally, human transport (Banks et al. 2015) and host naivety of the European chestnut could have contributed substantially to the rapid spread without the need for diversification and adaptation in the bridgehead population. As a response to fungal pathogen invasions among bats, some host populations evolved resistance (Langwig et al., 2017). There is no evidence for the emergence of tolerance or resistance in the European chestnut, however deployed control measures such as the artificial introduction of the *Cryphonectria hypovirus I* (CHV-1) can severely reduce fungal virulence (Rigling & Prospero, 2018). Interestingly, the mycoviral spread should be facilitated in asexual populations such as the invasive S12 lineage due to the lack of vegetative compatibility barriers. Outcrossing populations often harbor many different vegetative compatibility groups slowing transmission (Robin & Heiniger, 2001). Hence, the presence of the mycovirus may favor sexual reproduction and the immigration of additional vegetative compatibility types. In turn, the diversification may reduce the evolutionary advantage of the invasive lineage.

## Materials and Methods

### Samples of *C. parasitica*

A total of 188 virulent and mycovirus-free isolates of *C. parasitica* were sequenced. A majority of 182 originated from Albania, Bosnia, Bulgaria, Croatia, Georgia, Greece, Italy, Kosovo, Macedonia, Serbia, Slovenia, Switzerland, and Turkey (Fig. 1B, Supp. Table 1, Supp. Fig. 7). The six other isolates were from South Korea (*n*=2) and North America (*n=*4) (Supp. Table 1). A total of 125 European isolates belonged to the vegetative compatibility type EU-12, whereas 57 isolates represented other vegetative compatibility types (EU-types) occurring in central and south-eastern Europe. A total of 45 isolates from Bulgaria, Greece, Italy and Macedonia were already included in a previous population-wide study on south-eastern European *C. parasitica* diversity by (Milgroom et al., 2008). All samples were collected between 1951–2018 and are stored as glycerol stocks at –80°C in the culture collection of the Swiss Federal Research Institute WSL.

### DNA extraction and genotyping

All isolates were first inoculated onto cellophane-covered potato dextrose agar plates (PDA, 39 g/L; BD Becton, Dickinson & Company; Franklin Lakes, USA) (Hoegger et al., 2000) and incubated for a minimum of one week at 24°C, at a 14 h light and 10 h darkness cycle. After a sufficient amount of mycelium and spores had grown, the isolates were harvested by scratching the mycelial mass off the cellophane, transferring it into 2 ml tubes and freeze-drying it for 24 h. For DNA extraction, 15–20 mg of dried material was weighted and single tube extraction was performed using the DNeasy Plant Mini Kit (Qiagen, Hilden, Germany). DNA quantity was measured using the Invitrogen Qubit 3.0 Fluorometer (Thermo Fisher Sci-entific, Waltham (MA), USA) and DNA quality was assessed using the Nanodrop 1000 Spectrophotometer (Thermo Fisher Scientific, Waltham (MA), USA). Prior to sequencing, all isolates were screened for their genotype at 10 microsatellite markers (Prospero & Rigling, 2012). Additionally, the isolates were screened for their vegetative compatibility and mating type alleles in two multiplex PCRs, as described in Cornejo et al. (2019). Allele sizes for the genotyping and assays were scored with GeneMapper 5 (Thermo Fisher Scientific, Waltham (MA), USA).

### Illumina whole-genome sequencing, variant calling and filtration

Isolates were prepared for sequencing using the TrueSeq Nano DNA HT Library Preparation kit (Illumina, San Diego, USA). The libraries were sent to the Functional Genomics Centre Zurich (ETH Zurich and University of Zurich) and sequenced on the Illumina Hiseq 4000 platform (Illumina, San Diego, USA). The obtained sequences were trimmed with Trimmomatic v0.36 (Bolger et al., 2014) and aligned with Bowtie 2 v2.3.5.1 (Langmead & Salzberg, 2012) and SAMtools v1.9 (Li et al., 2009) to the *C. parasitica* reference genome (43.9 Mb) EP155 v2.0 available at the Joint Genome Institute (http://jgi.doe.gov/). Variant calling, selection and filtration were conducted with the Genome Analysis Toolkit GATK v3.8 and v4.0.2.0 (McKenna et al., 2010). We retained variants satisfying the following filtration parameters: QUAL: ≥ 100, MQRankSum (lower): ≥ -2.0, QD: ≥ 20.0, MQRankSum (upper): ≤ 2.0, MQ: ≥ 20.0, BaseQRankSum (lower): ≥ - 2.0, ReadPosRankSum (lower): ≥ -2.0, ReadPosRankSum (upper): ≤ 2.0, BaseQRankSum(upper): ≤ 2.0 (Supp. Fig. 1). Furthermore, we used BCFtools v1.9 (Narasimhan et al., 2016) and the R package vcfR (Knaus & Grünwald, 2017; R Core Team, 2014) for querying and VCFtools v0.1.16 (Danecek et al., 2011) for downstream variant filtering. Variants were additionally filtered for minor allele count (MAC) ≥ 1, excluding all missing data. Sites were also filtered per genotype, only keeping biallelic SNPs with a minimum depth of 3 and a genotyping quality (GQ) of 99. To exclude SNPs associated with the mating type, we ran an association study with TASSEL 5 (Bradbury et al., 2007) and retrieved *p*-values for each SNP across the genome. We set a *p*-value threshold to remove all SNPs with p ≤ 1×10^−10^ with the mating type for further analysis. The SNPs showing strong association with the mating type were located on scaffold 2 between positions 100’853–1’997’710 bp (Supp. Fig. 2).

### Phylogenetic reconstruction

The filtered whole-genome SNP dataset was used to build phylogenetic trees. We generated a maximum likelihood (ML) tree with RAxML v8.2.11 (Stamatakis, 2006), applying rapid bootstrapping generating 100 maximum likelihood (ML) trees. Phylogenetic trees were displayed using FigTree v1.4.3 (Rambaut, 2016). We also generated an unrooted phylogenetic network with using SplitsTree v4.14.6. (Huson & Bryant, 2006). SplitsTree was also used for calculating the PHI test (Bruen et al., 2006) to test for recombination. The required file conversions for using RAxML and SplitsTree (*i*.*e*. from VCF to FASTA format) were done with PGDSpider v2.1.1.5 (Lischer & Excoffier, 2011). We also performed a principal component analysis (PCA) as implemented in the R package ade4 (Bougeard & Dray, 2018).

### Inference of S12 donor populations

We generated an averaged co-ancestry matrix as inferred by fineSTRUCTURE v2.1.3 (Lawson et al., 2012). The software uses a Markov-Chain-Monte-Carlo (MCMC) based algorithm to infer ancestral contributions based on patterns of haplotype similarity. We ran the fineSTRUCTURE pipeline in ‘automatic mode’, with 500 Expectation-Maximation (EM) and 300’000 MCMC iterations, 400’000 maximization steps to infer the best tree and with ploidy set to 1. fineSTRUCTURE input files were created with LDhat (McVean & Auton, 2007) and the R package vcfR.

### Population genetic analyses and SNP impact assessment

We computed allele frequencies and estimated the allele spectrum using VCFtools. We used RStudio (RStudio Team, 2015) and ggplot2 (Wickham, 2016) for visualizations. Synonymous and non-synonymous sites were identified and annotated using SnpEff v4.3t (Cingolani et al., 2012). Variants which were classified by SnpEff as having a “high”, “moderate” or “modifying” impact on fungal protein sequences, were further processed in R using the packages dplyr (Wickham et al., 2018), reshape2 (Wickham, 2007), tidyr (Wickham & Henry, 2018), as well as and ggplot2. Furthermore, we performed a genome-wide association study (GWAS) with TASSEL 5 for identifying highly associated SNPs with genetic groups such as S12. Nucleotide diversity was calculated with vcftools --site-pi function using vcf-files that were converted to diploid. Identification of putative selective sweeps was performed using RAiSD v2.5 software that is based on three different genetic signatures of positive selection (with -y 1 -M 0 -w 50 -c 1 parameters) (Alachiotis & Pavlidis, 2018). The Manhattan plot shows the distribution of the computed composite *µ* statistic of positive selection.

### Transposable element *de novo* identification and population scans

We performed a *de novo* identification of transposable elements in the *C. parasitica* reference genome EP155 v2.0 using RepeatModeler v2.0.1 (Flynn et al., 2019). The consensus sequences were merged to the RepBase (RepBaseRepeatMaskerEdition-20181026) and then used for repeat annotation using RepeatMasker v4.0.7 with a blast cutoff at 250 (Smit et al., 2015). Repeats were then filtered out for low complexity and simple repeats, and parsed using the parseRM_merge_interrupted.pl script from https://github.com/4ureliek/Parsing-RepeatMasker-Outputs. We only retained elements longer than 100 bp and overlapping identical elements were merged into single elements for the annotation. In addition, elements of the same family separated by less than 200 bp were considered as part of the same TE and merged into a single element. We analyzed population-level presence/absence variation of transposable elements using the R-based tool ngs_te_mapper, using the previously trimmed reads and bwa version 0.7.17-r1188 (Bergman, 2012; Li & Durbin, 2009; Linheiro & Bergman, 2012). Overlapping genes and TEs were identified using the intersect function from the bedtools suite v2.29 (Quinlan & Hall, 2010).

### Genome compartmentalization and copy-number variation analyses

We investigated the genome architecture of *C. parasitica* following the protocol described in (Saunders et al., 2014). Briefly, we computed intergenic distances with the Calculate_FIR_length.pl script using the gene prediction for *C. parasitica* reference genome EP155 v2.0. We defined 40 bins given the range of 3’ and 5’ intergenic distances and calculated the number of genes that fell within each bin. To infer copy-number variation, we computed the normalized read depth for all isolates using the CNVcaller pipeline (Wang et al., 2017). Briefly, the reference genome was first split into 800 bp overlapping kmers that were re-aligned to the reference using blasr (-m 5 --noSplitSubreads --minMatch 15 –maxMatch 20 --advanceHalf --advanceExactMatches 10 --fastMaxInterval --fastSDP --aggressiveIntervalCut -- bestn 10) to identify duplicated windows (python3 0.2.Kmer_Link.py ref.genome.kmer.aln 800 ref.genome.800.window.link). The resulting windows were then used to calculate the normalized read depth for each isolate from the aligned reads (bam format) in 800 bp windows (Individual.Process.sh - b $ bam -h $ { i% .bam} -d ref.genome.800.window.link -s scaffold_1). For all further analyses, we used the read depth normalized for the absolute copy number and GC content of each sample (RD_normalized output). Given the observed distribution of the normalised read depth (NRD) across all isolates, we considered windows with NRD > 1.6 to be duplicated and windows with NRD < 0.4 as deleted.

### Mating experiments

The ability of *C. parasitica* isolates belonging to S12 with mating type MAT-1 to sexually outcross with isolates of opposite mating type, was assessed in an inoculation experiment. For this, we randomly selected four S12 and one non-S12 isolate with mating type MAT-1, as well as five isolates of opposite mating type MAT-2 from populations in Italy (Nebrodi), Kosovo (Molliq) and Bosnia (Konic, Vrnograc, Projsa) (Supp. Table 2). All isolates of both mating types belonged to the vegetative compatibility type EU-12 and mating pairs were only formed between isolates from the same geographic source populations. As substrate for the pairings, 40 mm long segments of dormant chestnut (*C. sativa*) stems (15–20 mm in diameter) were split longitudinally in half. The wood pieces were autoclaved twice, placed onto sterile petri dishes (90mm diameter), which were filled with 1.5% water agar to enclose the pieces. Mating pairs were inoculated on opposite sides of each halved stem with three replicates per pairing. The inoculated plates were then incubated at 25°C under a 16 h photoperiod (2500 Lux) for 14 days. After two weeks, mating was stimulated by adding 5 ml sterile water to the plates to suspend and distribute the conidia produced by both isolates over the stem segment. Any excess water was subsequently removed and the plates were incubated at 18°C under an 8 h photoperiod (2500 Lux). After 5 months of incubation, perithecia formation was assessed under a dissecting microscope. To confirm successful outcrossing, single perithecia were carefully extracted from the stromata and crushed in a drop of sterile distilled water. The resulting ascospore suspensions were plated on PDA and incubated at 25°C for 24–36 h. Afterwards, single germinating ascospores were transferred to PDA and incubated in the dark for 3–5 days at 25°C. DNA was then extracted from 10 mg of lyophilized mycelium using the kit and instructions by KingFisher (Thermo Fisher Scientific, Waltham MA, USA). All single ascospore cultures were screened for mating types by performing a multiplex PCR following the protocol described in Cornejo et al. (2019).

## Acknowledgements

We are grateful to Ursula Oggenfuss for helpful comments on a previous manuscript version. We thank Sandrine Fattore, Hélène Blauenstein, Quirin Kupper, Eva Augustiny, Dario Rüegg and Silvia Kobel for laboratory assistance. We acknowledge the Genetic Diversity Centre (GDC), ETH Zurich, for technical support and facility access. Ludwig Beenken and Martin Wrann helped with documenting mating experiments. Paolo Cortesi, Michael Milgroom, Kiril Sotirovski, Mihajlo Risteski, Marin Ježić and Seçil Akilli kindly provided samples. We thank Pierre Gladieux and Nikhil Kumar Singh for discussions and sharing scripts for data analyses. We are grateful for insightful discussions with Daniel Rigling. Funding was awarded to SP by the Swiss National Science Foundation (grant 170188) and to DC by the Fondation Pierre Mercier pour la science.

## Notes

#### Summary of Updates

Added Figures 4+5 with new analyses. New co-author.

