## Supplementary Figure for "Emergence and diversification of a highly invasive chestnut pathogen lineage across south-eastern Europe"

### Supplementary Figures

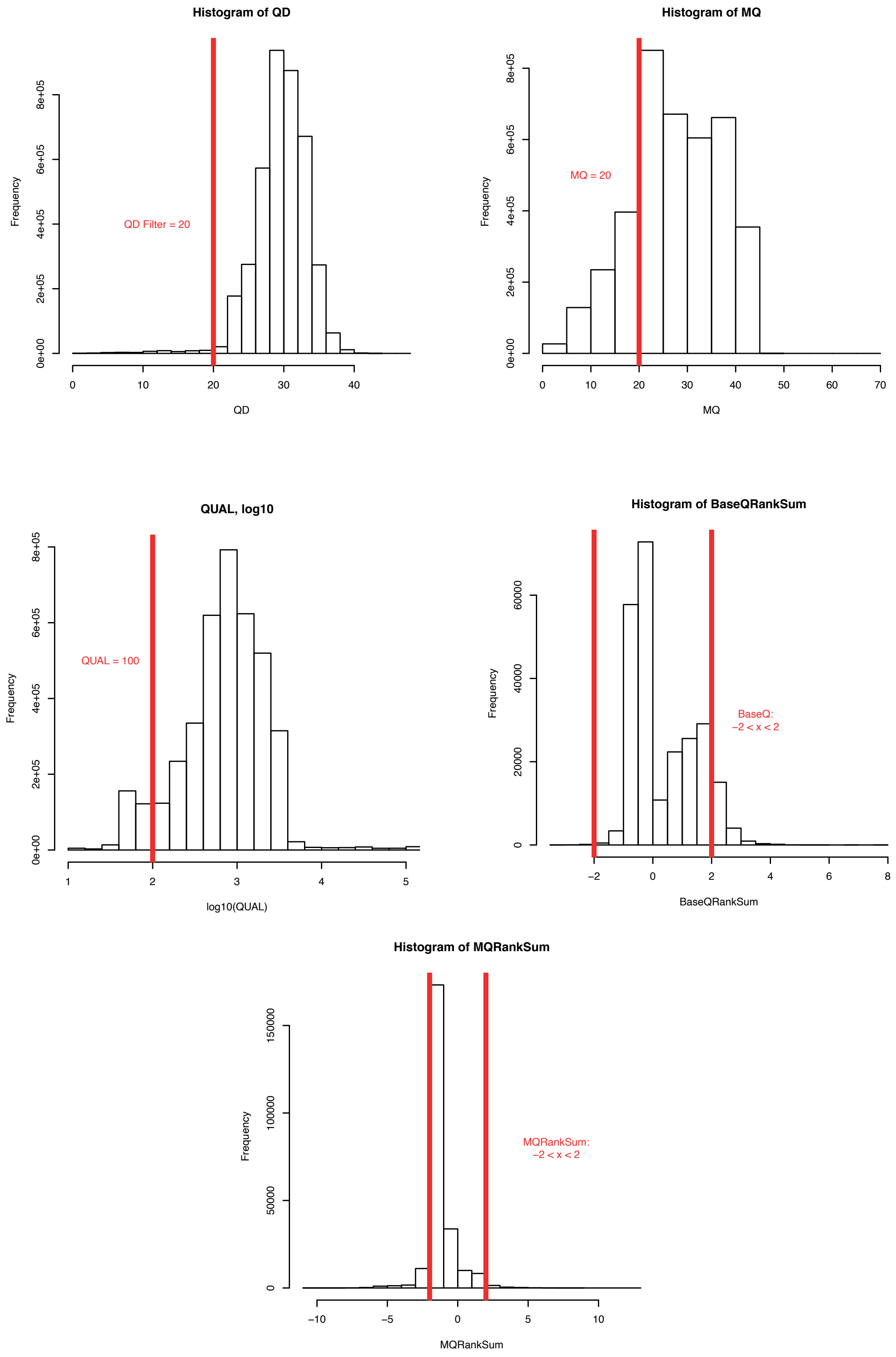


**Supplementary Figure 1:** Distribution of observed SNP filtration quality values.

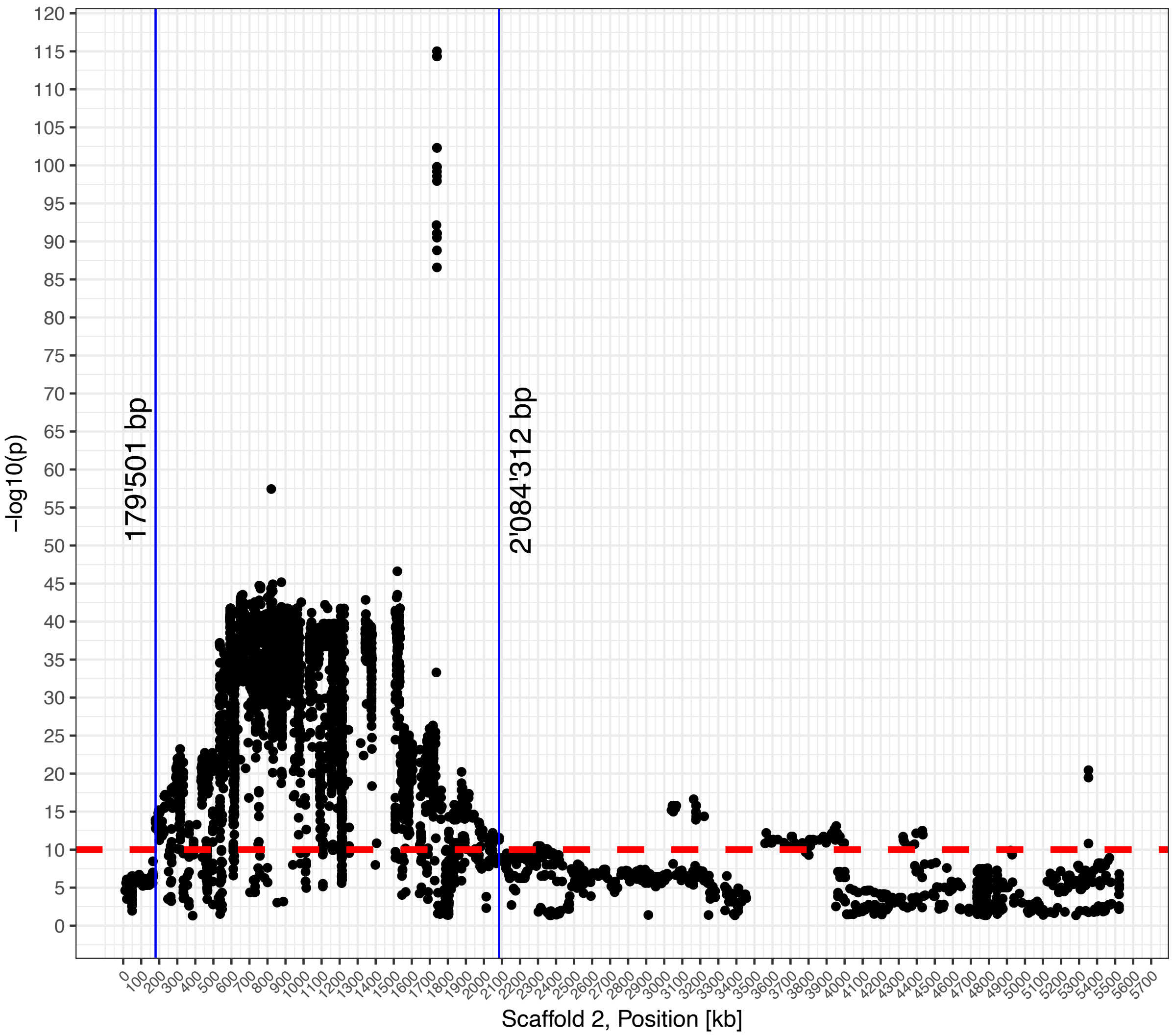


**Supplementary Figure 2**: Association mapping (GWAS) outcome for the mating type locus. *p*-value based threshold for removal of mating type locus (MAT) associated SNPs

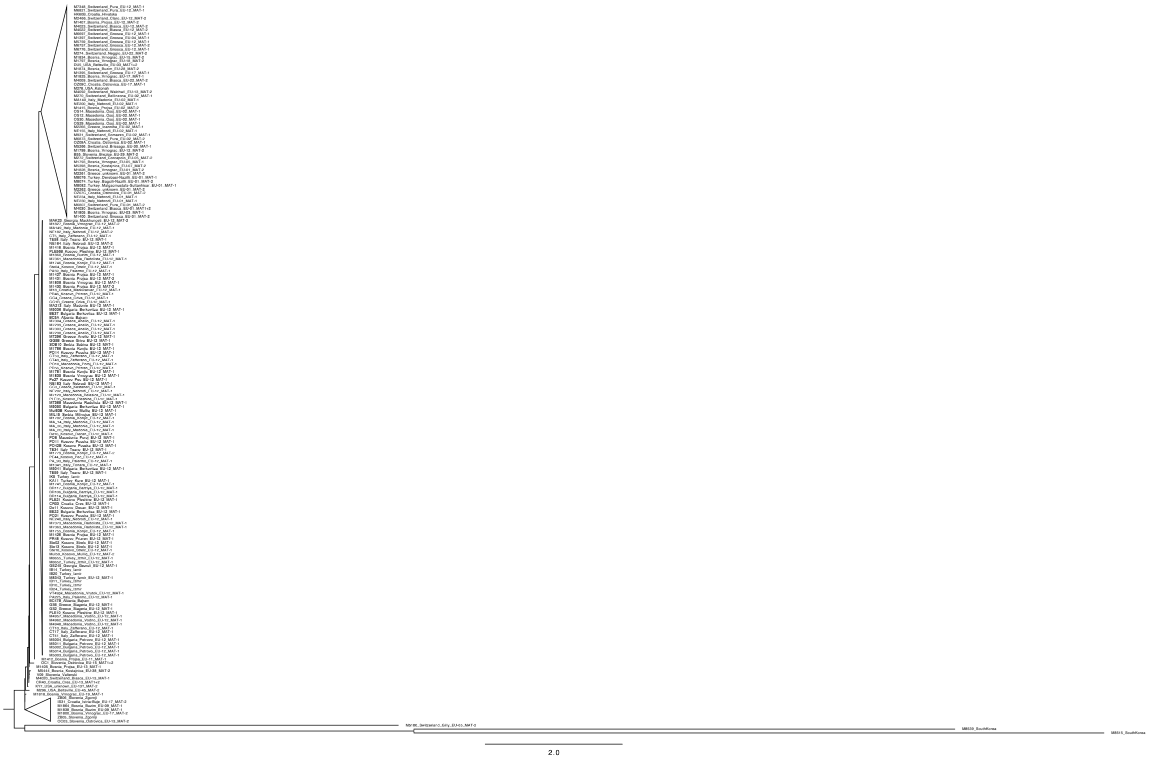


**Supplementary Figure 3**: Maximum-likelihood tree generated using RAxML showing full isolate identifiers

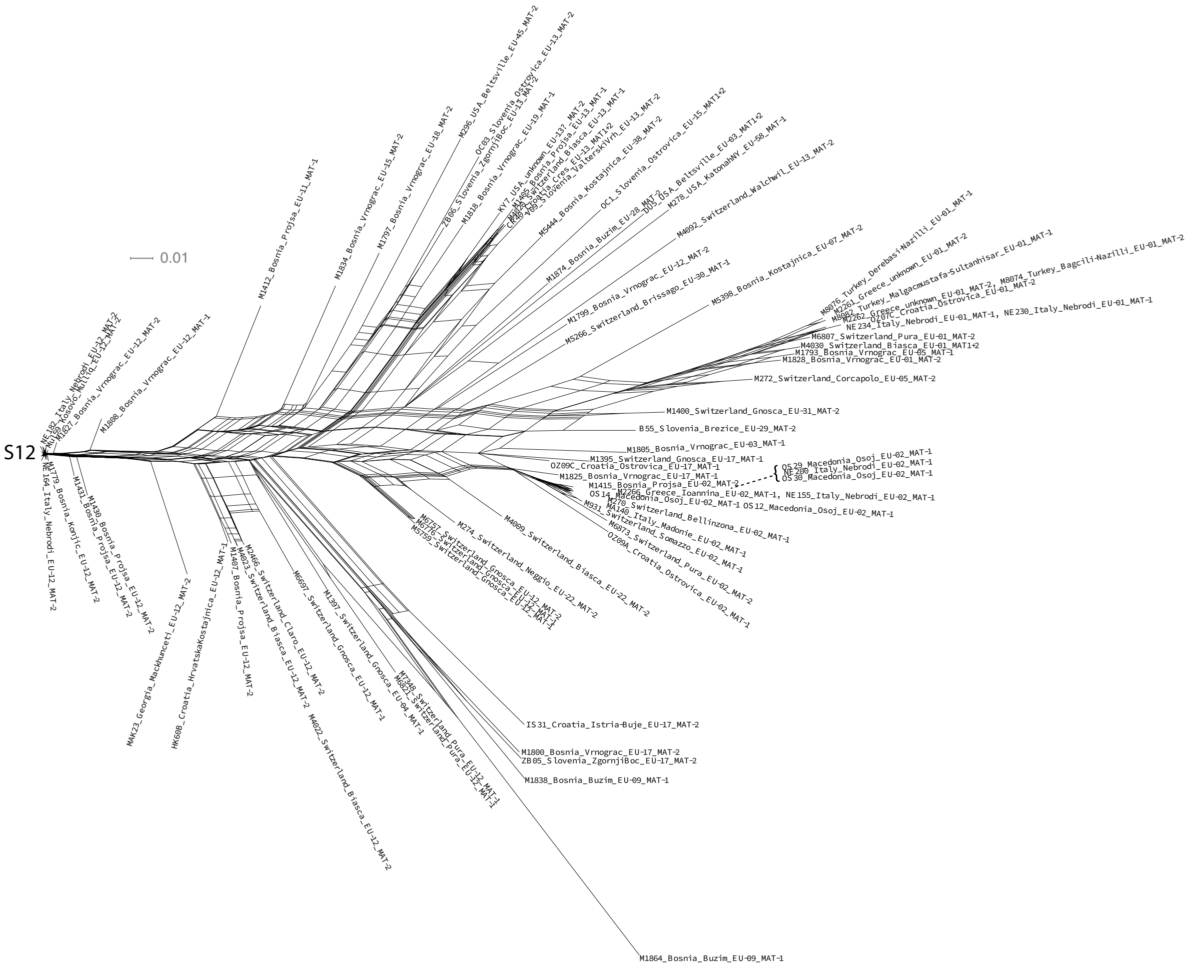


**Supplementary Figure 4:** Splitstree of all non-S12 isolates showing full isolate identifiers. Isolates identified as S12 are listed in Supplementary Table 1.

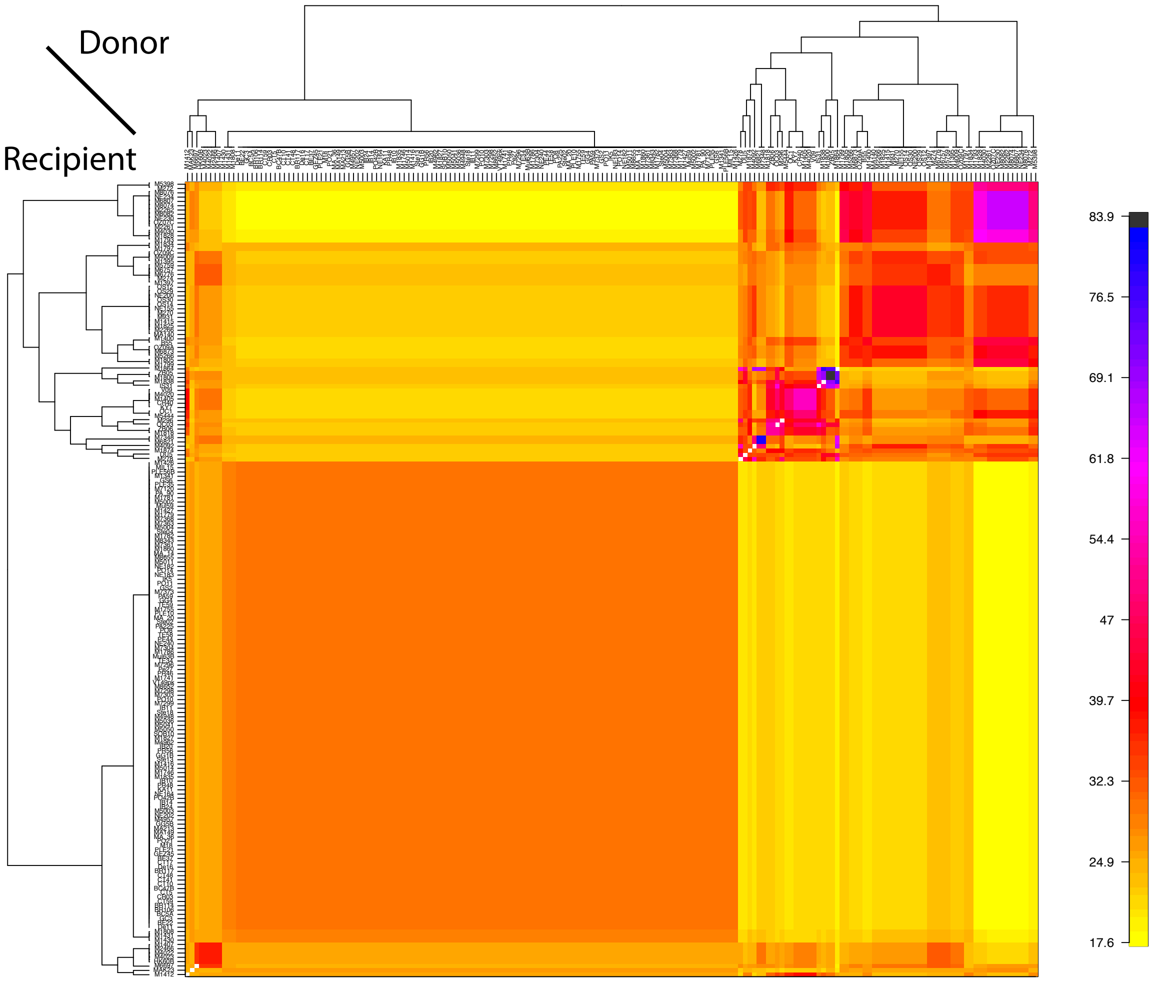


**Supplementary Figure 5:** Full averaged co-ancestry matrix showing full isolate identifiers.

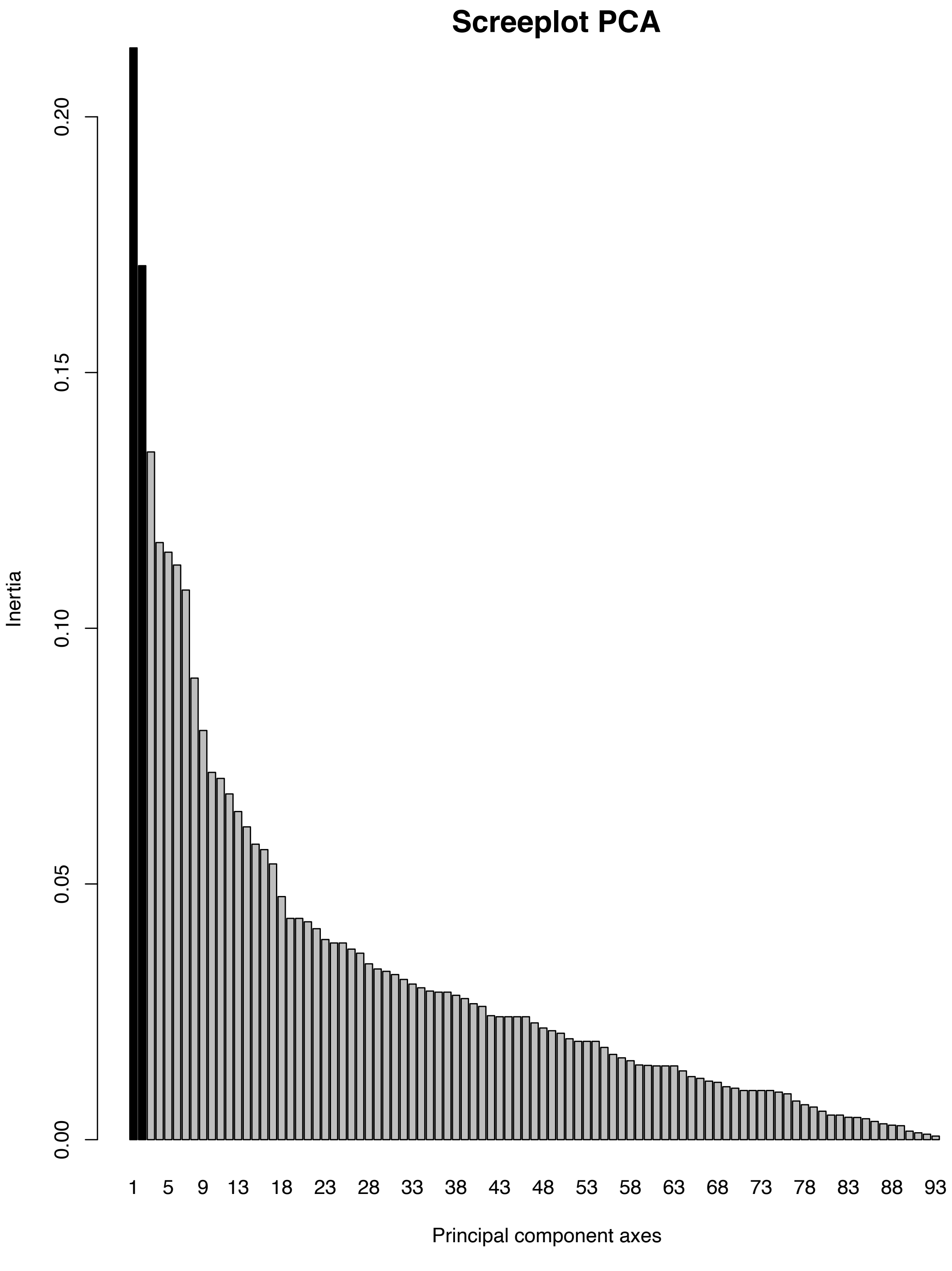


**Supplementary Figure 6:** Screeplot of the principal component analyis of the S12 lineage (see also Figure 4B).

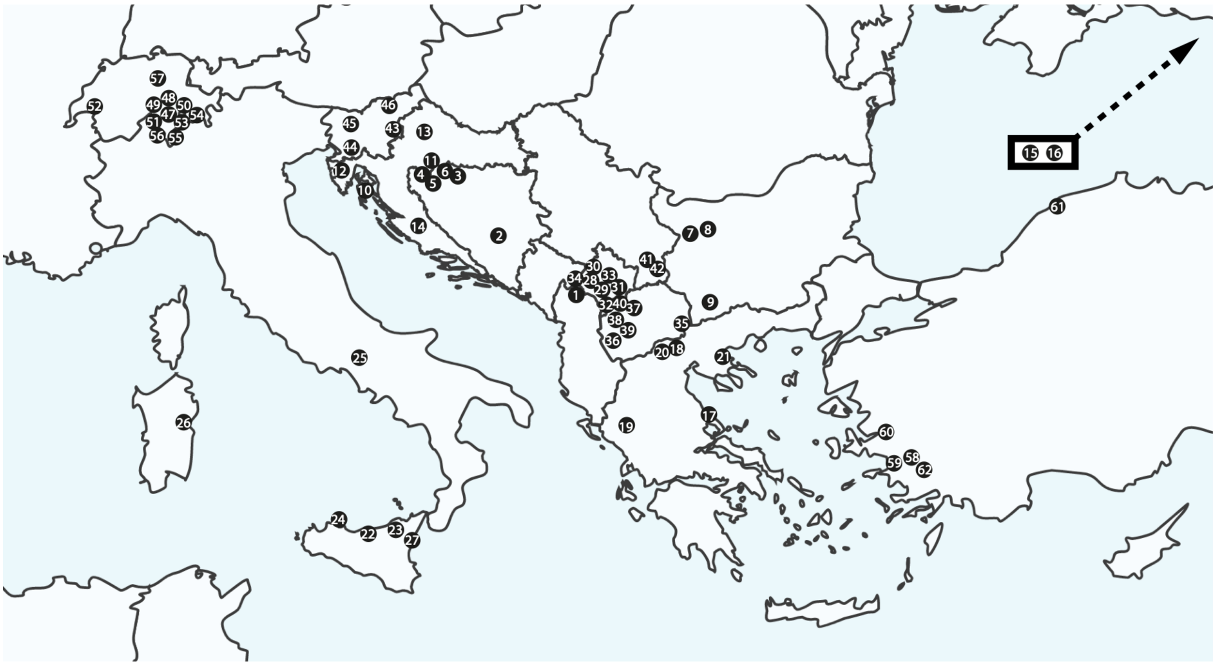


**Supplementary Figure 7:** Map of the European sampling locations, numbers correspond to sampling regions, as listed in Supplementary Table 1 (column "Europ.PopNr")
